# Portable Take-Home System Enables Proportional Control and High-Resolution Data Logging with a Multi-Degree-of-Freedom Bionic Arm

**DOI:** 10.1101/2020.05.19.102921

**Authors:** Mark Brinton, Elliott Barcikowski, Tyler Davis, Michael Paskett, Jacob George, Gregory Clark

## Abstract

This paper describes a portable, prosthetic control system for at-home use of an advanced bionic arm. The system uses a modified Kalman filter to provide 6 degree-of-freedom, real-time, proportional control. We describe (a) how the system trains motor control algorithms for use with an advanced bionic arm, and (b) the system’s ability to record an unprecedented and comprehensive dataset of EMG, hand positions and force sensor values. Intact participants and a transradial amputee used the system to perform activities-of-daily-living, including bi-manual tasks, in the lab and at home. This technology enables at-home dexterous bionic arm use, and provides a high-temporal resolution description of daily use—essential information to determine clinical relevance and improve future research for advanced bionic arms.

## 1 Introduction

Electromyography (EMG) from the residual forearm has been used to control commercially available and research-grade prosthetic arms (Jacob A. George et al., 2018; Hargrove et al., 2017; Kuiken et al., 2016; Mobius Bionics, 2020; Ottobock, 2017; Page et al., 2018; Perry et al., 2018; Touch Bionics Inc., 2017; Wendelken et al., 2017). Although research has demonstrated proportional control of multiple, independent degrees of freedom (DOFs) (Davis et al., 2016; Jacob A. George et al., 2018; Page et al., 2018), commercially available prostheses still suffer from a variety of limitations, including limited number of pre-determined grips, temporal delay due to sequential inputs used to select grips, fixed output force (e.g., from classifier algorithms), extensive training that lasts days to weeks, and non-intuitive methods of control (e.g., inertial measurement units (IMUs) on residual limb or feet) (Biddiss & Chau, 2007; Mobius Bionics, 2020; Ottobock, 2017; L. Resnik et al., 2017, 2019; L. J. Resnik, Acluche, Borgia, et al., 2018; Roche et al., 2014; Touch Bionics Inc., 2017).

Dexterous control of multiple DOFs, and the training associated with them, are not always amenable to deployment on portable systems with limited computational power. Only a few pattern-recognition (i.e., classifiers)(Kuiken et al., 2016; Mastinu et al., 2018; L. Resnik et al., 2017; Simon et al., 2019) or direct control algorithms have been studied at home (Pasquina et al., 2015; Simon et al., 2019). However, a Kalman filter (Wu et al., 2006), modified with non-linear, ad-hoc adjustments (Jacob A. George et al., 2019; J. G. Nieveen et al., 2020) can provide a computationally efficient approach (Jacob A. George, Radhakrishnan, et al., 2020) to proportionally and independently control multi-DOF prostheses. Proportional control algorithms enable realistic and life-like prosthetic control and can induce device embodiment in transradial amputees (Page et al., 2018).

High temporal resolution of the position and forces applied to the prosthesis is necessary to describe the interactive and refined movements made possible with proportionally controlled prostheses. These data are also necessary to describe key aspects of actual prosthesis use: revealing when objects were manipulated; whether movements were performed unilaterally or bilaterally (for bilateral amputees); which grasps were preferred; how often each DOF was used; and when new inter-digit collaborative movements were employed.

In a previous publication, we demonstrated at-home use of a portable, prosthetic control system that relied on a modified Kalman filter to provide six-DOF, real-time, proportional control (Jacob A. George et al., 2019). Here we describe the portable system and tasks completed at home in greater detail, including how the modified Kalman filter is trained and implemented on the portable system, as well as the system’s ability to record an unprecedented dataset of EMG, hand positions, and force sensor values. This technology constitutes an important step toward the commercialization of dexterous bionic arms by demonstrating at-home use and the ability to record prosthesis use with high temporal resolution.

## 2 Materials and Methods

### 2.1 Design considerations

A portable take-home system designed to research advanced bionic arms should meet several criteria for optimal performance and data collection: (a) the system must accurately and efficiently control the prosthesis; (b) training of the control algorithm must not be too long or burdensome to prevent its daily use—and thus should include the ability to quickly load a previously trained control algorithm; (c) high-temporal-resolution data should be stored automatically so that researchers can study at-home use without influencing the users with in-person observation; and (d) the system must be easy to use and allow the user to adjust control preferences.

#### 2.1.1 Accurate and efficient control

For accurate and efficient control, the system must be able to record EMG from the residual forearm, predict new kinematic positions, and send those positions to the prosthesis quickly with minimal or no perceived delay between the intention to move and the movement itself. Previous work in our lab has demonstrated responsive control of prostheses at update cycles of 33 ms (30 Hz) using a modified Kalman filter (Jacob A. George et al., 2018; Kluger et al., 2019; Page et al., 2018; Wendelken et al., 2017). The goal of this work was to implement these algorithms on a portable computer and provide position updates in under 33 ms.

#### 2.1.2 Fast training for daily use

For daily use, training of the control algorithm should be intuitive and fast. The time required to train a control algorithm includes data collection while the participant mimics preprogrammed movements of the prosthesis (Jacob A. George, Tully, et al., 2020), and training of the control algorithm itself (e.g., training the modified Kalman filter matrices). When training, or retraining, is required, it should be as fast as possible to minimize the setup time prior to use. Lengthy setup and training could make advanced prostheses burdensome to incorporate into daily life and prevent their acceptance among amputees. The system should also allow reloading of a previously trained control algorithm on demand.

#### 2.1.3 Comprehensive record of unsupervised arm use

A common approach to measure prosthesis use is to place IMUs on the prosthesis and record movement acceleration and angular velocity (Graczyk et al., 2018; Hargrove et al., 2017; L. Resnik et al., 2017; L. J. Resnik, Acluche, Borgia, et al., 2018). However, this approach fails to discriminate between gross movements from the residual limb and actual movement of the prosthesis’s hand and wrist. Video collection via body cameras can be used to record actual prosthesis use and other metrics (such as compensation strategies), but require storage of large video files and time-intensive post-hoc analyses (Spiers et al., 2017). Furthermore, the presence of a video camera reminds study participants they are being watched even though lab personnel are not physically present. However, with a portable system, prosthesis use at home can be studied by recording every movement for each DOF. By also recording the force applied to DOFs, interactive prosthesis use can be discerned from passive arm movements, such as those that might occur during walking or exploratory hand movements that are not functionally directed. Beyond describing total prosthesis use, this rich dataset can reveal detailed, refined movements and collaborative interactions between DOFs— including the force applied with each movement.

#### 2.1.4 User-friendly control adjustments

Finally, a prosthetic control system should be easy to use and allow adjustments to fit unique preferences. This includes a quick and simple approach to turn the system on, train the control algorithm and load a previously trained control algorithm. Control adjustments could also include flexibility to lock a DOF during extremely dexterous tasks (which may be easier with select DOFs stationary), or to operate the prosthesis in a position-control or a velocity-control mode.

### 2.2 Hardware and signal acquisition

The components of the portable system are shown in Figure 1. The DEKA LUKE Arm (DEKA; Manchester NH, USA) has 6 DOFs including D1 adduction/abduction; D1 flexion/extension; D2 flexion/extension; coupled D3-D5 flexion/extension; wrist flexion/extension, which also includes a slight radial and ulnar deviation, respectively; and wrist pronation/supination. It also has 19 sensors: six that report the position of each DOF and 13 that report the forces on each digit—including four directions on D1, two on the D2, and one on each of D3, D4 and D5—and on the lateral, dorsal and palmar (distal and proximal) aspects of the hand. The prosthesis itself records the aggregated use (i.e., time) within bins of movement velocity and electrical current draw for each DOF. It also records the total time each sensor experienced various forces (ten bins from zero to a max of 25.5 N) and the total time each DOF spent in various positions (ten bins across range of motion, which varies by DOF).

**Figure 1.**
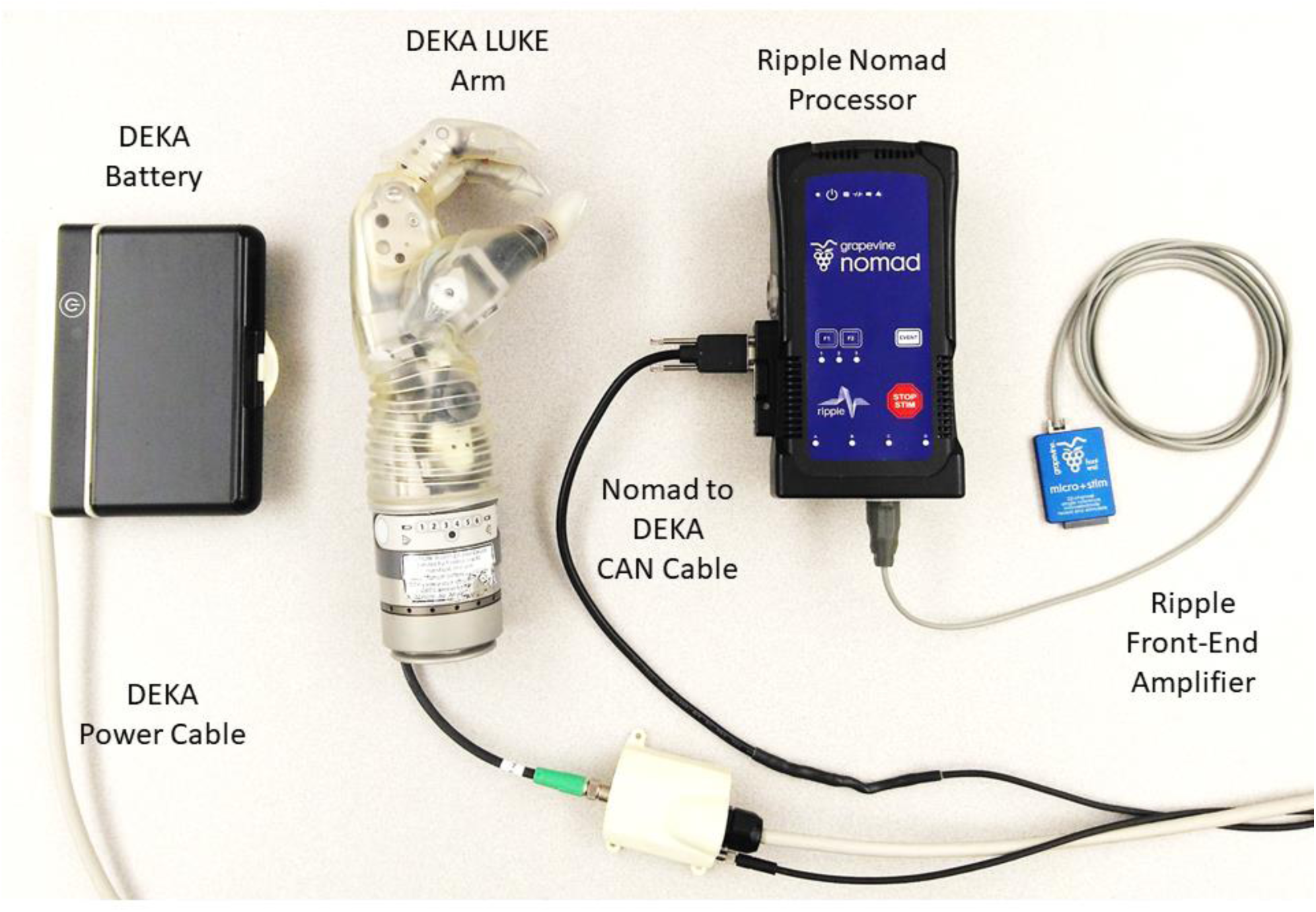
Portable take-home system for dexterous prosthetic control. The Ripple front-end acquires, filters, and amplifies EMG (at 1 kHz) to estimate motor intent using a modified Kalman filter with the battery powered Nomad neural interface processor. Communication occurs using a CAN protocol with the DEKA LUKE Arm to send commanded movements to the arm (at 30 Hz) and receive back the actual kinematic positions for six DOFs and the forces from 13 sensors (nine torque sensors, four pressure sensors at 30 Hz).

For the prosthetic control algorithm and data storage we used the Nomad neural interface processor (Ripple Neuro; Salt Lake City, UT, USA) for several reasons: an external, exchangeable battery provides up to four hours of power; wireless communication to external devices; 500 GB of hard disk storage; and up to 512 channels for data acquisition and stimulation. Ripple also provided firmware with the Nomad so that compiled prosthetic control algorithms can directly: acquire, filter and store EMG (1 kHz); communicate with and store data from the prosthesis (33Hz, CAN socket); start and stop via external buttons; and communicate over WiFi with external devices (TCP socket). Using a front-end amplifier (Figure 1; Ripple Neuro, Salt Lake City, UT, USA) we filtered (15 to 375 Hz bandpass; 60/120/180 Hz notch) the implanted EMG (iEMG) or surface EMG (sEMG) at 1 kHz. sEMG in intact participants was recorded with a Micro + Stim front-end (Ripple Neuro, Salt Lake City, UT, USA), and iEMG in the amputee participant was recorded with an active gator front end (Ripple Neuro, Salt Lake City, UT, USA). The Nomad runs Linux 8 (jessie) environment with an Intel® Celeron(tm) processor (CPU N2930) at 1.83 GHz with 2-GB RAM. Algorithms were converted to C using MATLAB® Coder and compiled for stand-alone use on the portable Nomad.

### 2.3 EMG feature calculation and decoding of motor intent

Training the prosthetic control algorithm (i.e., modified Kalman filter (Jacob A. George et al., 2019)) first requires the user to mimic preprogrammed movements of the prosthesis as it cycles through several movement trials for each DOF (Jacob A. George, Tully, et al., 2020). Features were then calculated for each differential EMG pair (496 total pairs from 32 single-ended electrodes) by taking the mean-absolute value of a moving 300-ms window (Figure 2c). Using the kinematic positions and the EMG features, the portable computer recursively chose 48 optimal features using the Gram-Schmidt forward-selection algorithm (Efron et al., 2004; Hwang et al., 2014; J. Nieveen et al., 2017) and computed the modified Kalman filter matrices (Wu et al., 2006).

**Figure 2.**
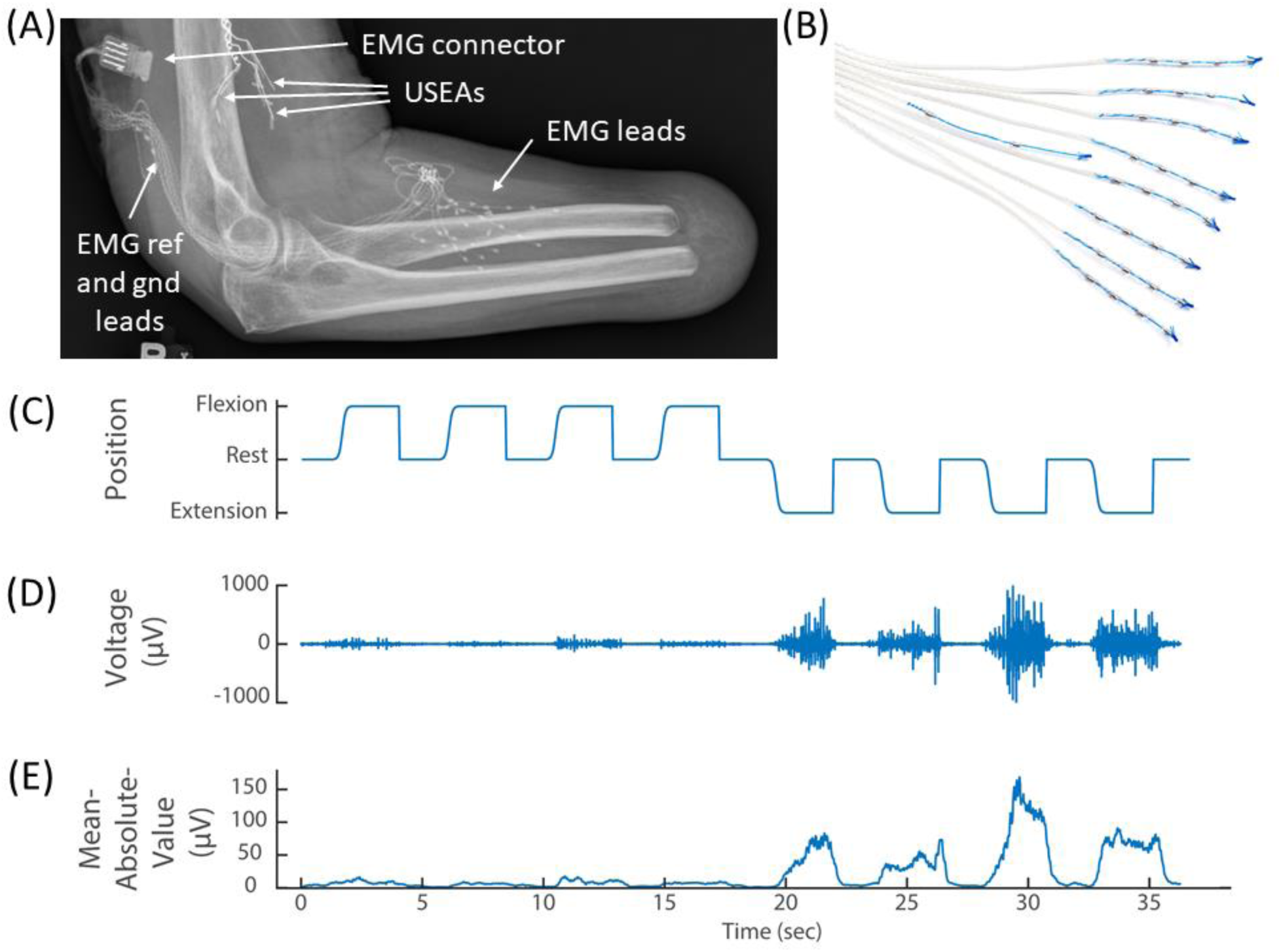
(A) X-ray of the elbow and residual forearm of a transradial amputee implanted with implanted three Utah Slanted Electrode Arrays (USEAs) and (B) 32 single-ended EMG leads (iEMG) and reference and ground. (C) The amputee mimics preprogrammed movements of the prosthesis (kinematics) while the portable Nomad system records iEMG voltage signals. (D) A representative iEMG channel that is active primarily during extension of the index finger. (E) A representative feature (mean absolute value of the iEMG channel in (C)) that is used to train the prosthetic control algorithm. Implanted Utah Slanted Electrode Arrays (USEAs) were not used with the portable system, but could be incorporated in future versions.

The Kalman filter presented by Wu et. al (Wu et al., 2006) was modified with external, ad-hoc thresholds as described in George et. al (Jacob A. George et al., 2019). To improve stability and reduce the effort required to sustain grasping movements, a latching filter has also added to the back-end of the modified Kalman filter (J. G. Nieveen et al., 2020).

### 2.4 Human subjects

Eight iEMG leads (Ripple Neuro; Salt Lake City, Utah, USA) with four electrodes each, and a ninth lead with an electrical reference and ground, were implanted in lower-arm extensor and flexor muscles as described previously (Jacob A. George et al., 2019)(Figure 2a-b). The electrode connector exited through a percutaneous incision and mated with an active gator connector (Figure 2a; Ripple Neuro; Salt Lake City, Utah, USA). This participant also had Utah Slanted Electrode Arrays implanted in the median and ulnar nerves but these devices were not used with the portable system. Surgical details have been previously described (Jacob A. George et al., 2019, 2018; Page et al., 2018; Wendelken et al., 2017).

Intact individuals were able to use the portable system with a 3D printed, custom-made bypass socket (Paskett et al., 2019) and a custom-made neoprene sleeve with 32 sEMG electrodes, plus one reference and one ground (Jacob A. George, Neibling, et al., 2020). As described previously (Paskett et al., 2019), the bypass socket is an open source device which suspends a prosthetic arm beneath the intact arm of the healthy volunteer and provides adequate range-of-motion so that the healthy volunteer can perform activities of daily living with an upper-limb prosthesis. The bypass socket was designed so that the electrode sleeve could be pulled up onto the forearm, locating the 32 recording electrodes over the extrinsic flexor and extensor hand muscles and the reference and ground electrodes over the ulna, about 2 cm distal to the elbow.

All experiments and procedures were performed with approval from the University of Utah Institutional Review Board.

## 3 Results

### 3.1 EMG recordings are consistent across desktop and portable systems

To ensure that the EMG was stored correctly on the Nomad, we concurrently recorded EMG with the portable system and a laboratory desktop system. As expected, concurrent recordings of sEMG on the portable and desktop systems were highly correlated (ρ = 0.95; p < 0.001) and the sEMG features were nearly identical (ρ = 0.99; p < 0.001). Due to slight variation in clock speeds, a temporal delay was observed (about 100 picoseconds/sample) which reduced the correlation coefficient. However, the correlation of the sEMG features suggests functional equivalency between the two recording systems.

### 3.2 Portable system offers a simple user interface and customizable control options

Three external buttons were employed to create a simple user-friendly interface. Pressing the first button initiated a new training session, which automatically granted control of the prosthesis to the user once training was complete (Figure 3a, see also Supplemental Video 1). The second button initiated a previously trained and compiled control algorithm (if available), so that the user could have on-demand control of the prosthesis. Finally, sequential inputs on a third button was used to toggle between position or velocity control modes or to freeze a DOF at a desired position.

**Figure 3.**
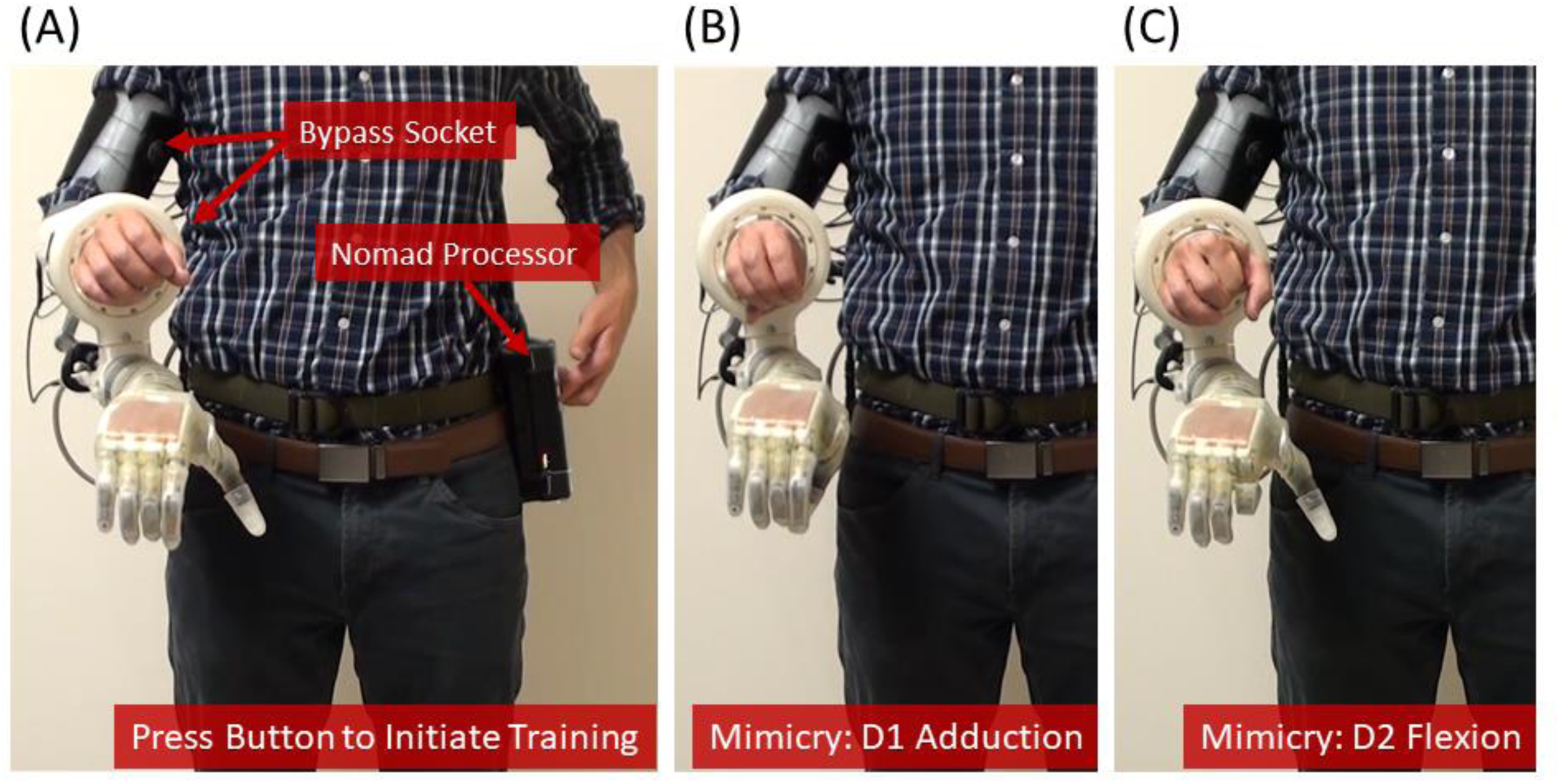
Training the prosthetic control algorithm with the portable system. (A) The user (shown here as an intact subject using a bypass socket *(Paskett et al. 2019)* to support the LUKE Arm) presses a button on the Nomad to start the training sequence, and then mimics the prosthesis while the Nomad cycles through each of six DOFs —(B) and (C) show D1 adduction and D2 flexion, respectively.

### 3.3 Portable system can be trained rapidly using steady-state modified Kalman filter

The system was trained in 7.5 minutes—including the time needed to collect the training data (252 sec) and the subsequent channel selection and computation of the modified Kalman filter matrices (about 199 sec) (Table 1). Training data included four trials of flexion and extension for D1, D2, D3/D4/D5, and the wrist; D1 adduction and abduction; wrist pronation and supination; and grasping and extending all digits together (Figure 3B-C, see also Supplementary Video 1). The trained modified Kalman filter was automatically saved to a log file and could be recompiled onto the Nomad as a stand-alone application for on-demand use (e.g., the second external button). This was accomplished over the Nomad’s wireless network using a laptop and required less than 30 seconds.

**Table 1:** Computational times required for training and testing (running) the steady-state, modified Kalman filter on the portable system.

| Process | Computation Time |
| --- | --- |
| *Training:* |  |
| Data collection | 252 s |
| Channel Selection | 198 s |
| Train Steady State Kalman Filter | 0.7 s |
| *Total Time* | 7.5 min |
| ***Testing:*** |  |
| Update Positions | 0.7 ms |

Prior to use, the steady-state modified Kalman gain matrix (K) was calculated by iteratively running the filter until the fluctuations in each value of the gain matrix were less than 1×10^-6^, reaching steady state after about 25 ms. With the gain (K), the observation (*H*) and the state-transition (*A*) matrices, a steady state matrix (Γ) was then calculated:

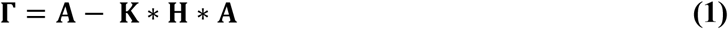

Thus, new position predictions 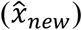 were calculated with only two matrix multiplications involving the previous positions 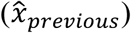 and 48 EMG features (z):

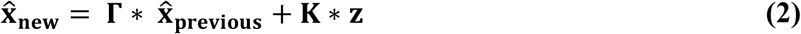

This simplification avoided a computationally expensive matrix inversion required by the recursive algorithm. Consequently, the time required to predict new positions and update the prosthesis was on average less than 1 ms, far below the update loop speed of 33 ms (Table 1).

### 3.4 Portable system can be used at home to complete various activities of daily living

The portable system has been used by intact participants to perform arm dexterity tests and activities of daily living in the lab (Figure 4), as well as to perform two-handed tasks at home (Figure 5). A transradial amputee also used the system at home, under staff supervision, to perform tasks of his choosing, some of which were not possible with his commercial prosthesis (Table 2 and Figure 6).

**Figure 4.**
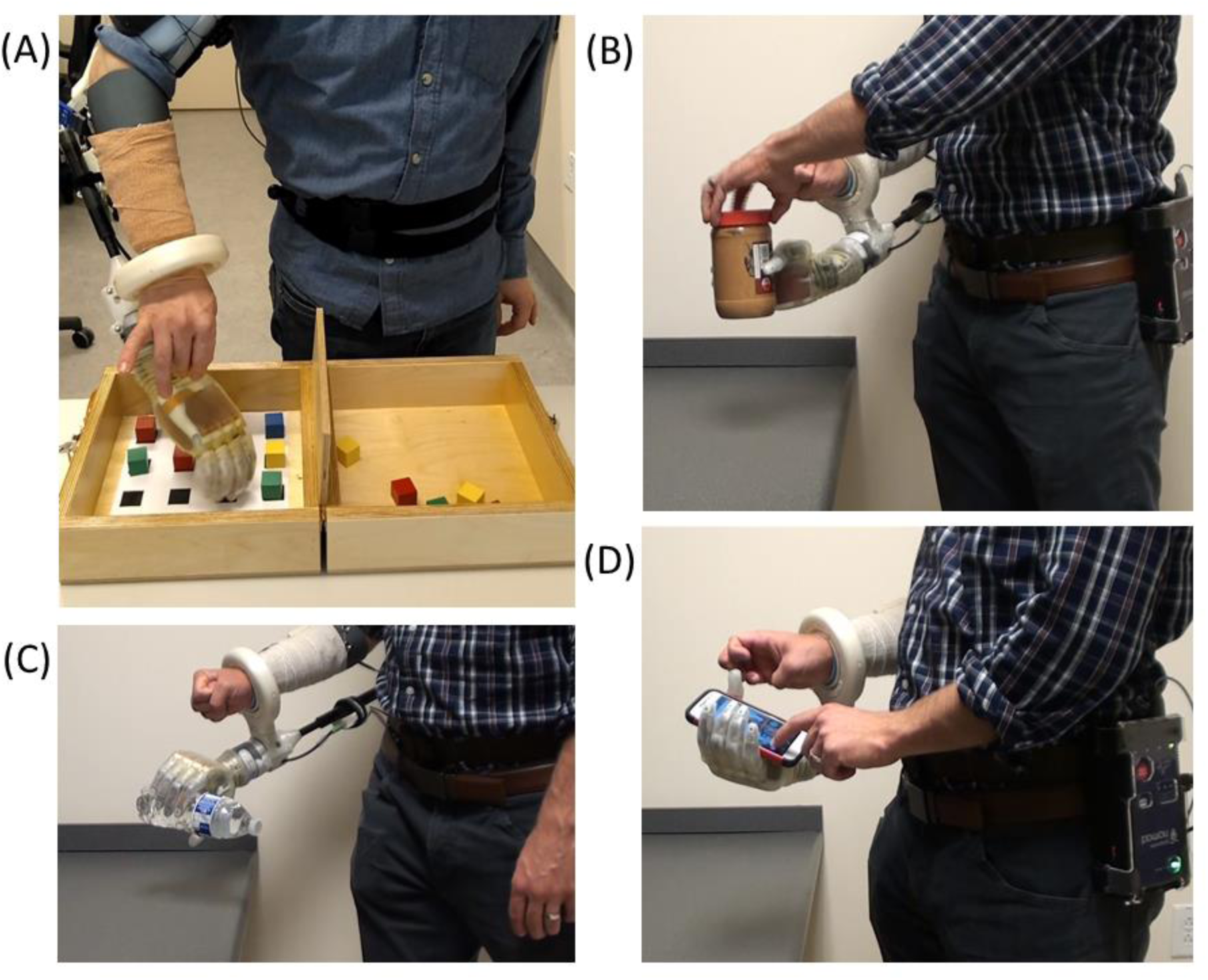
After the motor control algorithm was trained, intact participants used the portable system with a bypass socket in the lab to perform (A) an arm dexterity test and activities of daily living: (B) opening a jar; (C) pouring motion; and (D) using a smart phone.

**Figure 5.**
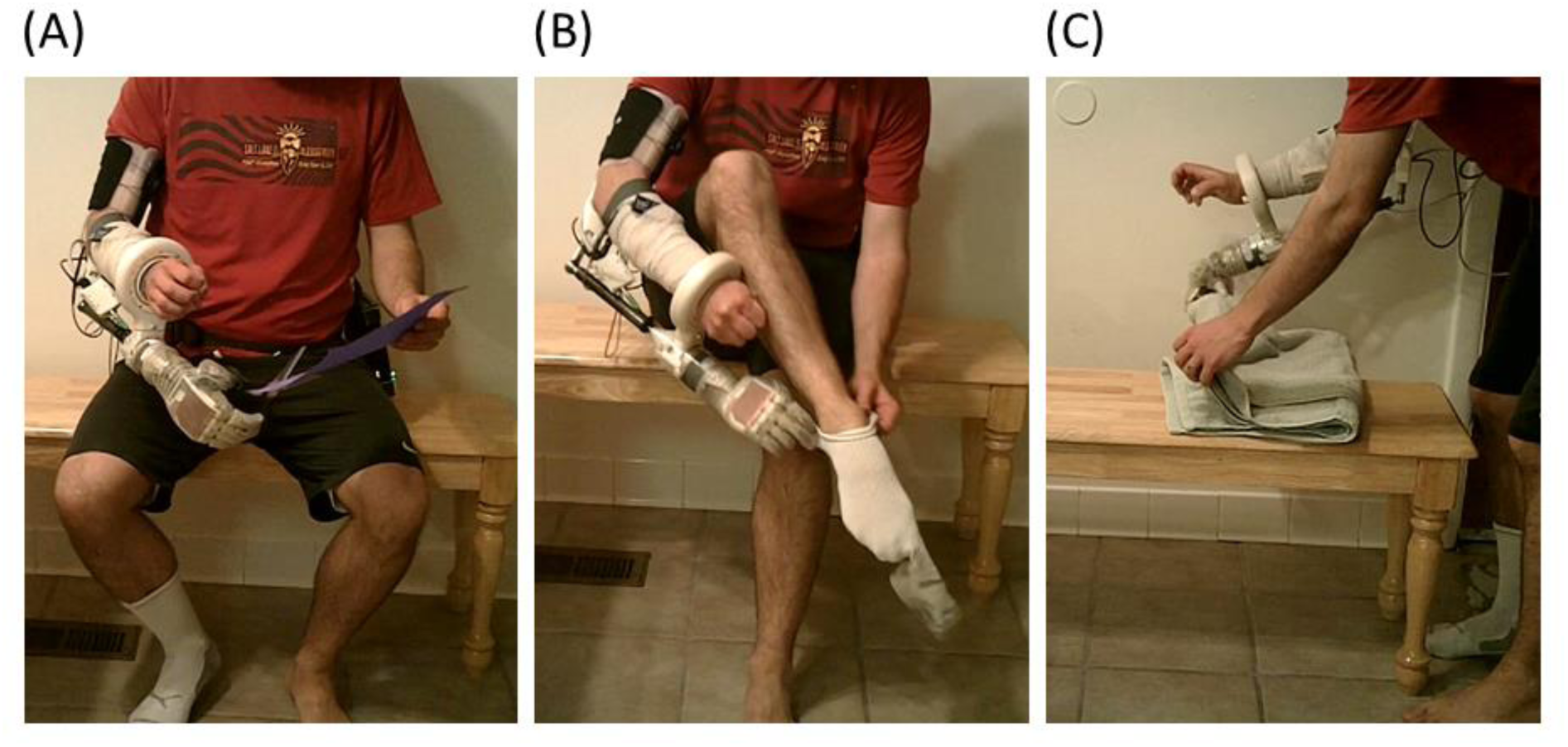
Two-handed activities of daily living at home using a bypass socket and the portable system: (A) using scissors; (B) donning a sock; and (C) folding a towel.

**Figure 6.**
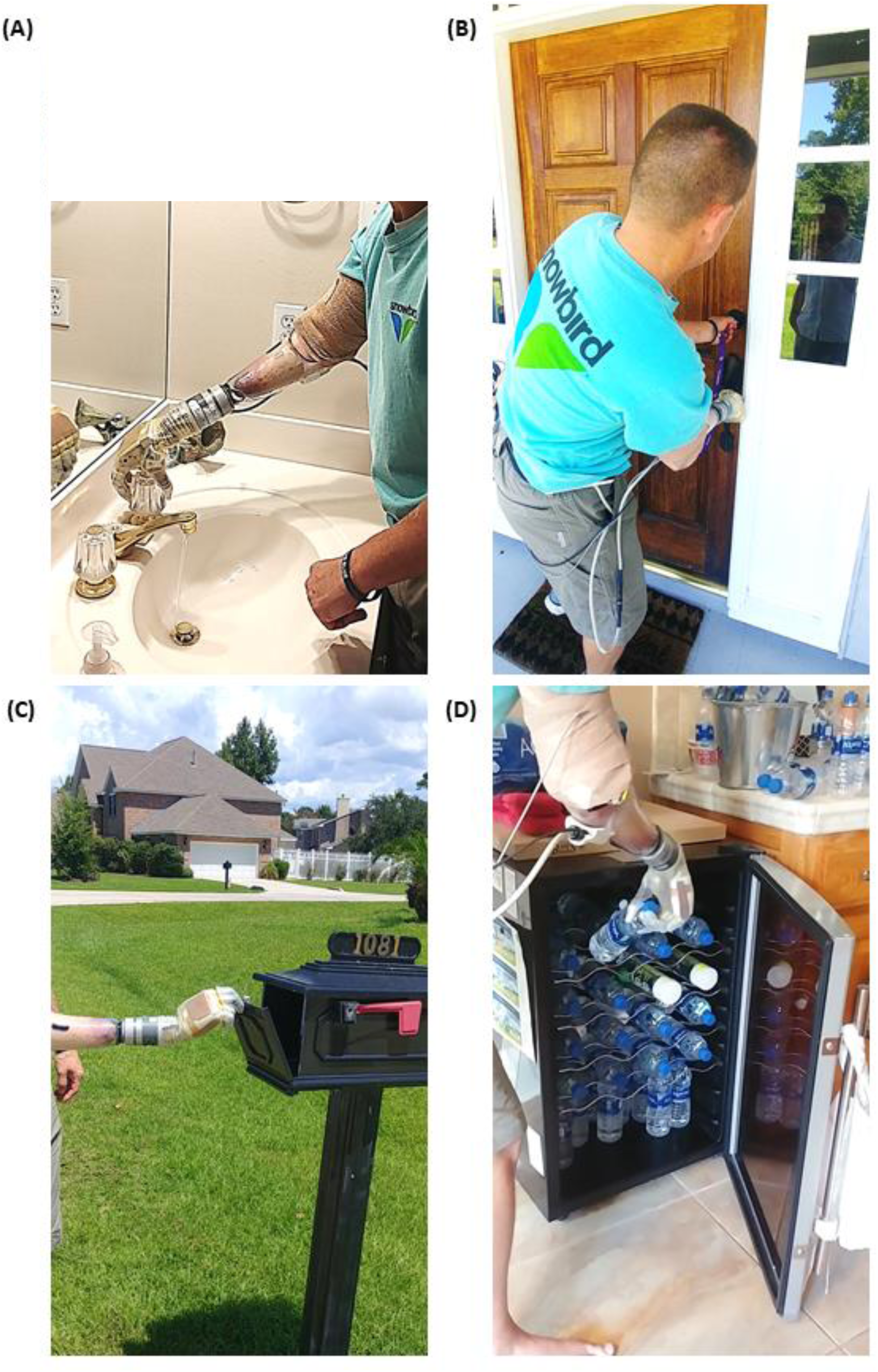
Transradial amputee performed supervised activities of daily living, of his own choice, at home using the portable take-home system. Images show the participant (A) turning faucet in the bathroom; (B) locking the dead-bolt on the front door; a bi-manual task not possible with his commercial prosthesis; (C) opening the mail box; and (D) retrieving water from the refrigerator.

### 3.5 Rich dataset from portable system reveals novel information about prosthesis use

EMG (sampled at 1 kHz), kinematic positions and forces applied to the prosthesis (both sampled at 30 Hz) were stored on the Nomad while a transradial amputee grasped, held and released an orange (Figure 7; see also Supplemental Video 4). Three phases of movement were clearly identified: preparing to grasp (when the index finger is near full extension); grasping (where the algorithm predicted the finger to be near full flexion but the orange restricted the actual position to about the rest position, which resulted in a dramatic increase in force); and releasing the orange (where the finger extended toward near full extension).

**Figure 7.**
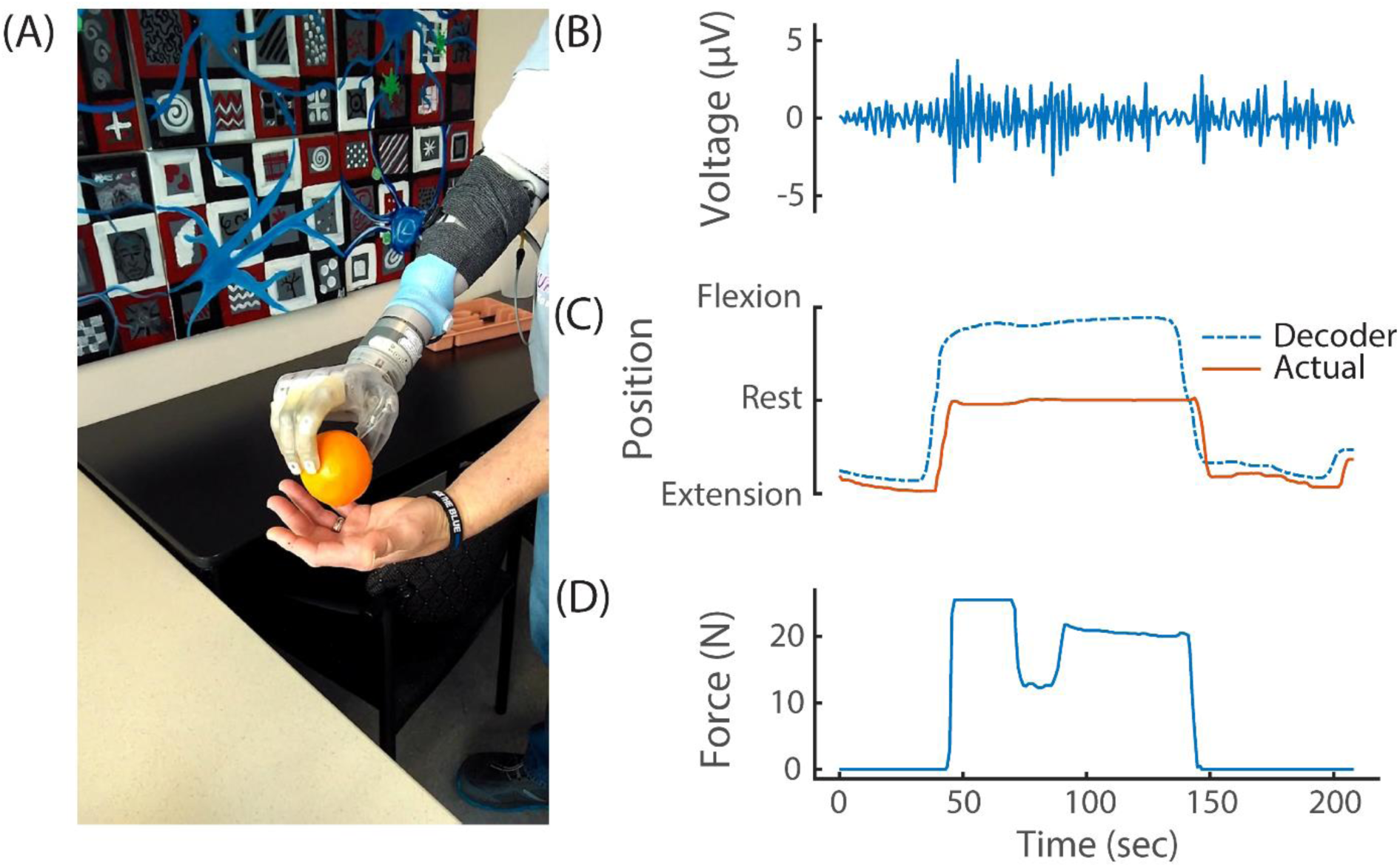
(A) Transradial amputee picking up an orange using implanted EMG electrodes and the portable system. During this task, the portable system recorded and stored (B) differential EMG (at 1 kHz), (C) kinematic output of the modified Kalman filter and actual kinematics of the prosthesis (at 30 Hz) and (D) prosthesis sensor values (at 30 Hz). For simplicity, only one differential EMG channel (of 48 total) and only one sensor (D1 pressure sensor; out of 13 total pressure and torque sensors and six DOFs) are shown.

Data are saved at a rate of 250 MB/hour in an ‘.hd5’ format. As a result, the 500 GB capacity of the Nomad can record nearly 2000 hours of arm use.

## 4 Discussion

We have shown that a modified Kalman filter can be trained in 7.5 minutes to proportionally control six independent DOFs using the Nomad, a portable electrophysiological recording system equipped with an ordinary processor. In addition, the time needed to acquire EMG and compute and update the prosthetic arm positions was less than 1 ms on average, far below the 33 ms update cycle, and provided real-time movement updates for the users.

The portable system also stores EMG, position and force data with unprecedented temporal resolution. This comprehensive dataset will be crucial for fully understanding how proportional control algorithms are used during unsupervised at-home use. Because of the Nomad’s large storage capacity and USB and Bluetooth connections it could also be configured to collect and store other types of data (e.g., video, bilateral arm use with IMUs).

To study at-home prosthetic use, previous take-home systems have stored limited usage data, including the time the device was turned on (Graczyk et al., 2018; Simon et al., 2019), aggregated hand movement (Simon et al., 2019), how often specific predefined grasps were used (Hargrove et al., 2017; Kuiken et al., 2016; Simon et al., 2019) or force applied to a limited number of sensors on the hand (Graczyk et al., 2018). Although these approaches may be sufficient for less refined control algorithms, to fully understand how proportional control is used, both high-temporal-resolution kinematic and force data for each DOF are necessary.

The example in Figure 7 highlights how the comprehensive data recorded reveals complex interactions between the various DOFs with proportional control. The stable D2 kinematics implies that the amputee held the orange with a fixed grasp from pick up to release; however, the force data revealed a dip in force during this same period. Close inspection of the kinematics from the opposing D1 also shows that a subtle readjustment occurred to improve the grasp stability (this can be seen in Supplemental Video 4). These refined movements are possible because of proportional control algorithms. Because DOFs are coupled together during object manipulation, the connection between each DOF must be considered.

Rich datasets like this will help researchers and clinicians study at-home, unsupervised use; improve prosthetic control algorithms and training paradigms (Jacob A. George et al., 2018) by understanding the types of grasps and DOFs commonly used; understand when mastery of prosthesis control occurs and when interventions might be applied or lifted; better describe noise encountered in real-world environments and design features and algorithms that reduce its influence on motor performance; and address many other unanswered questions about at-home use of advanced upper-limb prostheses. These rich datasets will also enable future at-home trials to study the benefits and use of high-resolution sensory feedback from intraneural electrical stimulation—a feature soon to be added to the portable system.

In contrast, previous data collection during at-home prosthetic use has relied on subjective surveys, usage logs, IMUs, and the amount of time the device is turned on to describe prosthetic use (Graczyk et al., 2018; Hargrove et al., 2017; L. Resnik et al., 2017; L. J. Resnik, Acluche, Borgia, et al., 2018). However, these approaches only approximate actual prosthesis use and could be misinterpreted. Some pattern recognition studies have recorded kinematic output and use of predefined grasps (Kuiken et al., 2016; Simon et al., 2019).

An important aspect of the portable system is the fast computation of position updates using a steady-state Kalman filter. We initially implemented the full recursive Kalman filter within each update cycle. However, the time required to complete the update was near and often exceeded our 33-ms update loop speed. Updating movement positions with the steady-state Kalman filter was quick (less than our loop time) and straightforward to implement. Our fast position update speeds will allow additional features to be added, including high-resolution, biomimetic, sensory feedback from intraneural (J. A. George et al., 2019; Wendelken et al., 2017) or electrocutaneous (Jacob A. George, Brinton, et al., 2020) stimulation.

The most computationally demanding aspect of training was performing Gram-Schmidt forward selection to choose the 48 most useful features out of the 496 differential pairs. Despite taking considerable time up front, this down-selection method has several advantages (J. Nieveen et al., 2017). First, choosing the features up-front enables fast loop speeds (below 33 ms) by eliminating the need to calculate complex features (e.g., principal components) or even all 496 differential EMG features during each update cycle. Second, forward selection recursively chooses independent and informative features using orthogonality reduction and correlation with the training kinematics. This ensures that each selected feature describes kinematics and not uncorrelated noise. Refined movements, the hallmark of proportional control algorithms, account for little variance and could be inadvertently discarded using techniques agnostic to the training kinematics. Finally, orthogonalization in the forward selection algorithm avoids redundant features and singularities.

Importantly, within eight minutes of powering the system on, the user can have real-time proportional control of six DOFs. The amount of time required to both collect training data by mimicking preprogrammed movements and to train the prosthetic control algorithm are related to the number of trials for each mimicked movement. In this work, and published elsewhere (Jacob A. George et al., 2019), an amputee familiar with the training process trained with only four trials on each DOF and a grasp and extension of all digits. With this training, he was able to control the prosthesis in the lab and perform tasks not possible with his commercial prosthesis at home (Jacob A. George et al., 2019). A less experienced user may require training with more trials; however, even if a naïve user requires twice as many trials, the total training time is still under 15 minutes.

An important question we have yet to fully explore is how often will retraining be required? The amputee in this study successfully used a trained algorithm the following day, suggesting daily training may not be necessary. It is also reasonable to believe that training with data collected across multiple days could provide better control (Jacob A. George, Neibling, et al., 2020), especially considering that iEMG leads are relatively stationary when implanted into residual musculature. However, training daily provides new users with the best control algorithm given the EMG features collected on that day and our experience suggests that over time they will become more stereotyped and thus have improved control. From the start, our training requires less time than pattern recognition algorithms which can require 14-40 hours of upfront, in-lab training with experienced professionals (L. Resnik et al., 2017; L. J. Resnik, Acluche, & Lieberman Klinger, 2018).

Currently, to use a previously trained control algorithm, the modified Kalman filter’s parameters must be recompiled into a stand-alone application on the Nomad. Although this process is very fast (less than 30 sec) and wireless, it currently requires an external computer running MATLAB®. Planned future work includes the ability to directly load trained parameters from a local file stored on the Nomad for on-demand use.

Ultimately, the ability to communicate with an application running on the Nomad was limited to one button (the other two could only start and stop a compiled application). Thus, we implemented sequential button pressing to selectively lock an individual DOF. The amputee used this feature to lock wrist rotation when using the system at home. For this amputee, poor wrist control was not uncommon and was likely due to dystonic muscle activity, common among those afflicted with complex regional pain syndrome and multi-year arm disuse prior to amputation (Jacob A. George et al., 2018). However, despite the amputee’s having low-amplitude EMG signals (e.g., Figure 7b), the modified Kalman filter algorithm provided control for five degrees of freedoms. Intact users did not lock any DOFs.

Future improvements will include wireless communication to a tablet or phone app where control selections can be easily made, communicated and saved locally on the Nomad. This will enable real-time adjustments including setting specific DOFs to velocity mode; adjusting the ad-hoc gains and thresholds of the modified Kalman filter on a DOF-by-DOF basis; and reloading a previous training or retraining the prosthesis with a modified training protocol if the first training was not satisfactory.

In its current form, the portable system is programmed to communicate only with the DEKA LUKE Arm. However, other custom communication sockets could be designed to communicate through the micro D-sub, USB or Bluetooth connections available to Nomad for proportional control of and data logging from other prosthetic limbs.

The importance of reliability in a take-home system cannot be understated—software and hardware must function as intended in the everyday environment. To fully test reliability, the system must be used at home, over many days and for many uses. To date, the system has been used on numerous occasions on campus and in the lab, but only at home by our laboratory staff and by one transradial amputee under staff observation (Jacob A. George et al., 2019). Ultimately, this system will be used in upcoming take-home clinical trials to record high-resolution data and study advanced, proportional control algorithms for upper-limb prosthesis use.

## 6 Author Contributions

MB, EB and TD designed the software running on the portable system. MB, TD, MP and JG tested the system with the human participants. MB wrote the manuscript. All authors revised the manuscript. GC oversaw all aspects of this research.

## 8 Acknowledgments

We thank Ripple Neuro, LLC for their generous support with custom firmware that enabled many features including high-resolution data collection and storage on the Nomad. We acknowledge Dr. Douglas Hutchinson and Dr. Christopher Duncan for their clinical support with the amputee participant.

